# HSP-1-Specific Nanobodies Alter Chaperone Function *in vitro* and *in vivo*

**DOI:** 10.1101/2025.08.25.672099

**Authors:** Nicholas D. Urban, Kunal Gharat, Zachary J. Mattiola, Ashley Scheutzow, Adam Klaiss, Sarah Tabler, Asa W. Huffaker, Monique Grootveld, Mary E. Skinner, Janine Kirstein, Matthias C. Truttmann

## Abstract

Targeted regulation of 70 kilodalton Heat Shock Protein (HSP70) chaperones, particularly the essential cognate heat shock protein (HSC70) and its *Caenorhabditis elegans* ortholog, HSP-1, may hold the key to improving cellular proteostasis and ameliorating aging-associated conditions linked to protein misfolding and aggregation. However, tools to selectively modulate HSP70 chaperone activity remain elusive. In this study, we pioneer the development of two novel nanobodies, B12 and H5, which specifically bind to both recombinant and endogenous HSP-1. We show that these nanobodies, differing by only two amino acids in their complementarity-determining regions, bind specifically to HSP-1 and effectively reduce both HSP-1 ATPase activity and protein folding capacity in a dose-dependent manner *in vitro*. We further demonstrate *in vivo* expression of B12, but not H5, in transgenic *C. elegans* strains reduces heat-stress survival and proteotoxic-stress resistance, mirroring the effects of *hsp-1* knockdown via RNA interference. Our findings suggest that these nanobodies can serve as effective and specific tools for modulating HSP-1 chaperone activity *in vivo*. These discoveries provide a foundation for future research exploring the therapeutic potential of HSP70-targeting nanobodies in aging and protein misfolding diseases.

## Introduction

The 70 kilodalton Heat Shock Protein (HSP70) family consists of conserved, ATP-dependent molecular chaperones critical for maintaining cellular proteostasis during stress^1^. Each monomer includes an approximately 44 kDa nucleotide-binding domain (NBD), which mediates ATP binding and hydrolysis, and an approximately 28 kDa substrate-binding domain (SBD) that contains a hydrophobic pocket for binding polypeptides^2,3^. The lid of the SBD regulates substrate interaction, with conformational changes initiated in the NBD through ATP hydrolysis, affecting the trapping of substrates in SBD^4–6^. This allosteric cycle transitions HSP70 between high- and low-affinity states for client engagement^7^. Targeted interactions with co-chaperones enhance HSP70 activity and provide for greater diversification in substrate interaction. J-domain proteins interact with the NBD of HSP70s to stimulate ATP hydrolysis, while nucleotide exchange factors (NEFs) interact with the NBD to facilitate ADP-ATP exchange, acting as molecular adaptors in physiological contexts^8–10^.

Functionally, HSP70 chaperones aid in the folding of newly synthesized proteins, prevent protein aggregation by stabilizing protein folding intermediates, refold and disaggregate misfolded and aggregated proteins, direct misfolded proteins to degradation pathways, and enable protein trafficking^6^. Dysregulated HSP70 function is implicated in diseases such as Alzheimer’s disease, Parkinson’s disease, and cancer metastasis^11–15^. With current clinical research exploring inhibitors and activators, HSP70s remain promising therapeutic targets for age-related diseases and conditions involving protein misfolding^16,17^.

The HSP70 family is highly conserved across species, highlighting their fundamental role in proteostasis^6,18^. In humans, there are 13 HSP70 genes, a figure mirrored in mice; *Caenorhabditis elegans* have about seven functional HSP70 genes, although broader classifications that include pseudogenes can raise this total to between 10 and 16^17,19^. Heat Shock Cognate 71 kDa Protein (also known as HSC70 or HSPA8) is the constitutively expressed HSP70 family chaperone in mammalian cells, supporting protein folding, complex assembly, and the refolding or degradation of misfolded proteins^20^. In the face of cellular stress, HSC70 can translocate to the nucleus to avert aggregation of heat-denatured nuclear proteins, aiding nuclear stability^21–24^. In *C. elegans*, the HSC70 ortholog, HSP-1, fulfills a similar function, facilitating development, promoting resistance to stress, and regulating proteostasis by ensuring proper nascent polypeptide folding and minimizing protein aggregation^25,26^. Both homologous proteins also intersect with proteolytic pathways to maintain cellular quality control; thus, the shared roles of HSC70 and HSP-1 underscore the high degree of evolutionary conservation within the HSP70 family, affirming their significance in safeguarding organismal health and fitness across different species^1,26–30^. However, specific tools to modulate HSC70 and HSP-1 activity remain elusive.

In this study, we identify a pair of nanobodies which specifically bind and modulate the activity of the cognate *C. elegans*’ chaperone, HSP-1. Nanobodies, also known as VHH domains, are single-domain antibody fragments derived from the heavy-chain-only antibodies found in camelids (e.g., camels, llamas, and alpacas) and sharks^31,32^. Consisting solely of the variable domain, they measure approximately 15 kDa, enabling them to bind epitopes often inaccessible to larger antibodies, including enzyme active sites^33,34^. Their high specificity and affinity, combined with excellent solubility and remarkable thermal and chemical stability, make them valuable tools in both research and therapeutic contexts^35,36^. Nanobodies can be efficiently produced in bacterial systems, such as *Escherichia coli*, as well as expressed in eukaryotic systems where they retain functionality^37,38^. Given their tissue-penetration capabilities and rapid clearance from the bloodstream, nanobodies see increasing use in biosensing, as Positron Emission Tomography (PET) tracers, and as targeting entities of PROTACs^39–42^.

Here, we present two nanobodies, B12 and H5, which differ in only 2 amino acids and bind specifically to both recombinant HSP-1 *in vitro* and endogenous HSP-1 *in vivo* in *C. elegans*. Using ATPase and protein refolding assays, we show both B12 and H5 inhibit the ATPase activity and refolding capability of HSP-1 in a dose-dependent manner. Furthermore, we find that moderate *in vivo* expression of B12 is sufficient to phenocopy reductions in survival and protein misfolding (Aβ_1-42_)-induced paralysis akin to *hsp-1* knockdown in *C. elegans*. Taken as a whole, these results establish B12 and H5 as two specific nanobodies to specifically regulate HSP-1-dependent processes *in vitro* and *in vivo*.

## Materials and Methods

### Nanobody purification and labeling with sortase

Following a previously described protocol^43^, nanobodies were expressed in the periplasm of *E. coli* cells, extracted following outer membrane rupture, and retrieved using Ni-NTA beads. Eluted nanobodies were dialyzed to remove excess imidazole. For labeling of nanobodies with biotin or TAMRA, we mixed 10 μg of sortase with a 20-fold molar excess of GGG-biotin or GGG-TAMRA and incubated the reaction at 4 °C overnight. The following day, we removed unconjugated dye and biotin molecules using P10 desalting columns and unconjugated nanobodies using Ni-NTA.

### Chaperone protein purification

All *C. elegans* chaperones were purified according to previously described protocols^44,45^ 100 ng of pSumo plasmids containing the chaperone sequences were transformed into *E. coli* BL21(DE3) (New England BioLabs, #C2527I). A starter culture was prepared by inoculating 30 mL of LB supplemented with Ampicillin (100 mg/ml) media with 5 transformants followed by incubation at 37 °C and shaking at 130 RPM overnight. The next morning, 2 L of LB-Ampicillin media were inoculated with 20 mL of the starter culture and incubated at 37 °C and shaking at 130 RPM. When OD_600_ reached 0.6, protein expression was induced by adding 1 mM IPTG and continuing incubation at 20 °C with shaking at 130 rpm overnight. The next morning, bacteria were pelleted by centrifugation at 6000 rpm for 30 minutes at 4 °C. Pellet was thawed in ice and resuspended in 100 mL of lysis buffer (30 mM HEPES pH 7.4, 500 mM KAc, 5 mM MgCl_2_, 20 mM imidazole, 10% glycerol, 1 mM PMSF, 1 mM β-mercaptoethanol, 10 µg/ml DNase I, 1 tab/50 ml of cOmplete EDTA-free protease inhibitor cocktail (Roche)). The cell suspension was sonicated (Branson 450 Sonifier) for 10 minutes (20 seconds on; 40 seconds off) at 50% amplitude and soluble fraction was recovered after centrifugation at 16,000 rpm (Sorvall RC6+, Thermo Scientific) for 30 min at 4 °C. In total, 3 mL of Ni-NTA slurry (High-Density Nickel 6BCL-NTANi, Agarose Bead Technologies) were added to the soluble fraction, and His-tagged protein binding was allowed to occur at 4 °C with gentle rotation over a period of 1 hour. Slurry was filtered through a gravity flow column and washed with 25 ml of high-salt (30 mM HEPES pH 7.4, 1 M Kac, 5 mM MgCl_2_, 20 mM imidazole, 10% glycerol, 1 mM β-mercaptoethanol) and low-salt (30 mM HEPES pH 7.4, 50 mM Kac, 5 mM MgCl_2_, 20 mM imidazole, 10% glycerol, 1 mM β-mercaptoethanol) buffers. His-tagged proteins were recovered after addition of 4 mL of elution buffer (30 mM HEPES pH 7.4, 50 mM Kac, 5 mM MgCl_2_, 300 mM imidazole, 10% glycerol, 1 mM β-mercaptoethanol) and incubation for 30 min at 4 °C with gentle rotation. Elution fraction was transferred to a 12-14 kDa MWCO dialysis membrane (Spectra/Por 2, Spectrum laboratories) and buffer was exchanged overnight at 4 °C against 2 L of dialysis buffer (30 mM HEPES pH 7.4, 50 mM Kac, 10% glycerol, 1 mM β-mercaptoethanol) supplemented with 100 µL of 0.76 mg/ml His-Ulp1 protease to cleave the His-Smt3 tag. To remove the cleaved tag and the protease, protein solution was incubated with 1.5 mL of Ni-NTA slurry for 30 min at 4 °C and filtered through a gravity flow column. The recovered protein was aliquoted and flash-frozen in liquid nitrogen for storage at −80 °C.

### Microscale thermophoresis (MST)

MST experiments were performed in duplicate using a Monolith NT.115 instrument (Nanotemper Technologies). HSP-1 was labeled with RED-NHS using the Protein Labeling Kit RED-NHS 2^nd^ Generation (#MO-L011) following the manufacturer’s protocol. For binding assays, 20 nM of labeled HSP-1 was mixed with a 16-step serial titration of each binding partner and incubated for 30 minutes in the dark prior to measurement. For HSP-1:H5 binding experiments, concentrations ranged from 0.0033 to 109.4 μM; for HSP-1:B12, concentrations ranged from 0.003815 to 125 μM; and for HSP-1:Enhancer, the concentrations ranged from 0.0106 to 350 μM. Data were analyzed using MO.Control v2.7.1 software. Binding constant (K_d_) was determined using the built-in K_d_ model, and the binding curve was normalized as Fraction Bound.

### Luciferase refolding assays

Luciferase assay was performed as previously described with slight modifications^46^. Briefly, a 3 nM luciferase solution in 1x dilution buffer (50 mM HEPES pH 7.4, 100 mM Kac, 5 mM MgCl_2_, 1 mM DTT, 10 µM BSA, 3.5 µM Pyruvate Kinase (Roche), 3 mM Phosphoenol Pyruvate) was denatured at 45 °C for 15 minutes. Next, luciferase was diluted to a final concentration of 1 nM in 1x dilution buffer containing 5 µM HSP-1, 0.25 µM HSP-110, and 5 µM DNJ-13 and amount of nanobody as described in the figure. After 2 hours, 5 µL aliquots from each refolding sample were dispensed to three different wells of a 96-wells white polystyrene plate (MultiScreen® 96-Well-Plate, Millipore) containing 100 µL of assay buffer (25 mM glycylglycine, 100 mM Kac, 15 mM MgCl_2_, 5 mM ATP). Then, 100 µL of 1 µM luciferin solution was added to each well, and luminescence was measured using a plate reader (Infinite 200 PRO, Tecan). Attenuation was not used, integration time was 1000 ms, and settle time was 0 ms. Values were normalized to the values measured for the Trimeric-chaperone complex sample after 2 hours and data were presented as the percentage of luciferase activity recovered after 2 hours.

### ATPase assays

ATPase assays were performed as previously descried with slight modifications^46^ Briefly, 50 µL samples were prepared containing 1x reaction buffer (50 mM HEPES pH 7.4, 100 mM Kac, 5 mM MgCl_2_, 0.017% Triton X-100), 5 µM HSP-1, 0.25 µM HSP-110, and 5 µM DNJ-13 and amount of nanobody as described in the figure. ATP was added to initiate the reaction at a final concentration of 2 mM, followed by incubation at 20 °C for 1 hour. 10 µL aliquots of phosphate standards and of each sample were transferred in triplicates to a 96-wells transparent microplate (Greiner), followed by 160 µL of green malachite reaction solution (2:1:3 dilution of 0.082% green malachite, 5.7% ammonium molybdate (in 6 M HCl) and water) and 20 µL of 34% sodium citrate. Absorbance at 650 nm was measured in a plate reader (Infinite 200 PRO, Tecan). To determine ATPase activity (%) for each sample, measurements were normalized to the activity of the Trimeric-chaperone complex. The corresponding free phosphate concentration in each well was calculated using the equation derived from the phosphate calibration curve.

### C. elegans strain preparation and maintenance

All worms were maintained at 15 °C on standard NGM plates spotted with OP50-1 *E. coli* for at least two generations without starving before being used for experiments. MTX265 (*mtmEx100[myo-2p::mCherry; hsp-16.48p:: B12::HA])* and MTX277 *mtmEx103[myo-2p::mCherry; hsp-16.48p:: H5::HA])* were generated by SUNY biotech by microinjecting 10 ng/µL of marker and 10 ng/µL of transgene plasmid DNA into the germline of young adult N2 worms. *mtmEx100* was integrated via UV-irradiation to generate MTX298 (*mtmIs20[myo-3p::mCherry; hsp-16.48p:: B12*]) and backcrossed 5x to WT animals prior to being used in experiments. MT24068 (*nEx2480[myo-3p::mCherry; hsp-16.48p:: VHH-7::HA]* was generated by microinjection and integrated MT24421 (*nIs775[myo-3p::mCherry; hsp-16.48p:: VHH-7::HA]*). GMC101 (*dvIs100 [unc-54p::A-beta-1-42::unc-54 3’-UTR + mtl-2p::GFP]*) was obtained from the Caenorhabditis Genetics Center (CGC). MTX307 (*mtmIs20; dvIs100*), MTX319 (*mtmEx103; dvIs100*), and MTX329 (*nIs775; dvIs100*) were generated by the Truttmann Lab. Wild type (N2, Bristol) animals obtained from CGC and were used as reference controls unless otherwise stated. All strains were backcrossed at least 3 times before being used in experiments.

### Gene knockdown via RNA interreference

RNA interference was performed as previously described^47^. Briefly, HT115 *E. coli* expressing siRNA against the target gene of interest was grown in a 5x overnight culture of LB media. The following day, the overnight culture was spun down and the pellet was resuspended in fresh 1x LB media (e.g 5 mL of overnight culture resuspended in 1 mL of fresh LB). The 1x culture was supplemented with 100 mg/mL carbenicillin (1:1000, antibiotic, GoldBio, Cat #C-103-25) and 1M isopropyl β-D-1-thiogalactopyranoside (IPTG) (1:200, Dot Scientific, Cat #DSI5600-25). The culture was then spotted on to nematode growth media (NGM) plates that were supplemented with 1M IPTG (1:1000), 100 mg/mL carbenicillin (1:1000) and 10 mg/mL nystatin (1:1000, antifungal, Dot Scientific, Cat #DSN82020-10). Plates were used immediately on the day of preparation or kept at 4 °C for no more than 2-3 days. For all experiments, worms were synchronized onto HT115 *E. coli* expressing siRNA against *pos-1*, which is a zinc-finger transcription factor required for embryonic development and sterilizes worms without affecting the hatched animal^47–52^. All siRNA-expressing *E. coli* were obtained from the Vidal Library^53^.

### Worm synchronization

Worms were synchronized via hypochlorite treatment as previously described^47^. Briefly, animals were washed 3x with sterile filtered M9 and treated with 1 mL of hypochlorite solution. Animals were incubated for 8-10 minutes with shaking at room temperature. Following bleaching, eggs were pelleted using centrifugation (2900 rcf) and washed 2x with sterile M9 before being plated.

### Worm lysis for protein biochemistry

Worms were lysed as previously described^47^. Briefly, worms were washed 3x with sterile M9 buffer and snap frozen in liquid nitrogen then stored at -80 °C. Samples were resuspended in ∼200-400 μL of sterile filtered worm lysis buffer (HEPES (20mM, 7.4pH), NaCl (20mM), MgCl_2_ (200mM), and Nonidet P-40 (0.5%)) spiked with protease inhibitor cocktail (Pierce^TM^ Protease and Phosphatase Inhibitor Mini Tablets, EDTA-free, ThermoFisher, Cat #A32961). Worms were transferred to reinforced tubes with a steel ball and lysed using a Qiagen TissueLyser III (7.5 minutes, 30 Hz). Lysate was cleared at 16,100 rcf (4 °C, 15 minutes) twice, transferring to a precooled tube after each transfer. The soluble fraction was collected, and protein lysate concentration was determined using the Pierce^TM^ BCA Protein Assay Kit (ThermoFisher, Cat #23227) following the manufacture’s instruction.

### Immunoblotting

Either 1 μg of recombinant purified protein or 10-20 μg of *C. elegans* worm lysate was added to 4x Laemmli Protein Sample Buffer (Bio-Rad, Cat #1610747) as described by the manufacturer, then boiled at 100 °C for 5 minutes. Samples were subjected to SDS-PAGE and subsequently transferred to a PVDF membrane using the Bio-Rad Trans-Blot^®^ Turbo System (Bio-Rad, Cat #1704150) and Trans-Blot^®^ Turbo RTA Transfer Kit, PVDF (Bio-Rad, Cat #1704272) following the manufacture’s instruction. Membranes were blocked for 1 hour at room temperature with gently rocking and probed with the appropriate antibody or nanobody (1 μg/mL) overnight at 4 °C with gentle rocking. Membranes were washed 3x with sterile 0.1% TBS-Tween20 (TBS-T) and incubated with appropriate secondary antibodies for 1 hour at room temperature while rocking. The membrane was then washed 3x with TBS-T. Supplemental Table 1 lists all blocking buffers and antibodies used in this study.. Chemiluminescent signal was observed using Prometheus Protein Biology ProSignal® Dura ECL Reagent (Prometheus Protein Biology Products, Genesee Scientific, Cat #20-301) following the manufactures instructions. Membranes were imaged using an Invitrogen iBright1500. If necessary, membrane stripping was done using *OneMinute^®^*Western Blot Stripping Buffer (GM Biosciences, Cat #GM6001) following the manufacturers instruction, washed vigorously with ddH_2_O, then rehydrated in 0.1% TBS-T for 15 minutes. Membranes were then treated as described above. Immunoblot quantification was done using Fiji (version 2.14).

### Immunoprecipitation assays

Worms were synchronized and lysed as described above. Magnetic protein G agarose beads were washed with 300 μL of worm lysis buffer and separated using a magnetic rack 3 times (Dynabeads™ Protein G for Immunoprecipitation, Invitrogen^TM^, Cat. #10003). The appropriate amount of worm lysate (∼500-1000 μg) was diluted to an equal volume of worm lysis buffer, then precleared for at least 1 hour at 4 °C while rotating using washed beads. Following clearing, an aliquot was saved to use as input control (5% of total immunoprecipitation). For immunoprecipitation using exogenous nanobody, 20 μL/sample of pre-conjugated streptavidin magnetic beads (Dynabeads™ MyOne™ Streptavidin C1, ThermoScientific^TM^, Cat. #65001) were washed 3 times and added to the pre-cleared lysate. For immunoprecipitation of nanobodies from *in vivo* expression, 20 μL/sample of pre-conjugated anti-HA magnetic beads (Pierce^TM^ Anti-HA Magenetic Beads, ThermoScientific^TM^, Cat. #88836) were washed 3 times and added to the pre-cleared lysate. Lysate/bead mixture was incubated over night at 4 °C with rocking. The following day, the protein-bead complex was isolated using a magnetic rack and washed 3x with cold worm lysis buffer. Following the final wash, the complex was resuspended in ∼15-30 uL of 4x Laemmli Protein Sample Buffer (Bio-Rad, Cat #1610747) as described by the manufacturer and boiled at 100 °C for 5 minutes. Beads were then isolated using a magnetic rack, and the supernatant was collected and subjected to SDS-PAGE and treated as described above (Immunoblotting).

### C. elegans mild heat stress survival assays

Worms were synchronized via hypochlorite treatment as previously described above. Eggs were spotted onto NGM/RNAi interference plates described in *RNA interference* above with HT115 *E. coli* expressing siRNA against *pos-1*. Worms were then placed in a 20 °C incubator. Day 1 adults were transferred to 60 mm IPTG-NGM plates spotted with fresh HT115 with siRNA targeting *pos-1* or *hsp-1* (∼40-50 worms per plate). Animals were then transferred to a 25 °C incubator. For heat shock survival assays, animals were transferred to a 37 °C incubator (induced) for 30 minutes. Plates were left for one hour to return to room temperature, then placed at 20 °C. Dead animals, as confirmed by lack of spontaneous or prodded movement, were removed from the plate and counted every other day until the completion of the experiment.

### Paralysis assays

Paralysis assays were performed as previously described^47^. Briefly, animals were synchronized via hypochlorite treatment, then maintained at 15 °C until day 1 of adulthood. Animals were then transferred to fresh 60 mm IPTG-NGM plates spotted with fresh HT115 with siRNA targeting *pos-1* or *hsp-1* (∼40-50 worms per plate), then placed at either 25 °C or 15 °C. Animals were considered paralyzed if unable to complete a full body movement spontaneously or when prodded. Paralyzed animals were counted and removed from the plate when scored every other day.

## Results

### Nanobodies B12 and H5 bind specifically to HSP-1 *in vitro*

To generate nanobodies specific for HSP-1, we immunized an alpaca with recombinant HSP-1 protein purified from *E. coli* over-expression cultures. We then isolated peripheral lymphocytes, extracted RNA, and amplified the variable domain of the heavy chain (VHH)-coding sequences to clone them into a phagemid library. Next, we performed phage display on immobilized HSP-1 to enrich for HSP-1-binding VHHs. We then tested 90 VHH clones in a crude ELISA setup, of which 13 showed strong binding. Sequencing of the binders revealed that the best performing VHHs represented two unique VHH sequences, which we named B12 and H5 (**Fig. 1A**). Interestingly, B12 and H5 differ by only two amino acids in CDR1 and have identical CDR2 and CDR3 domains (**Fig. 1A**).

**Figure 1.**
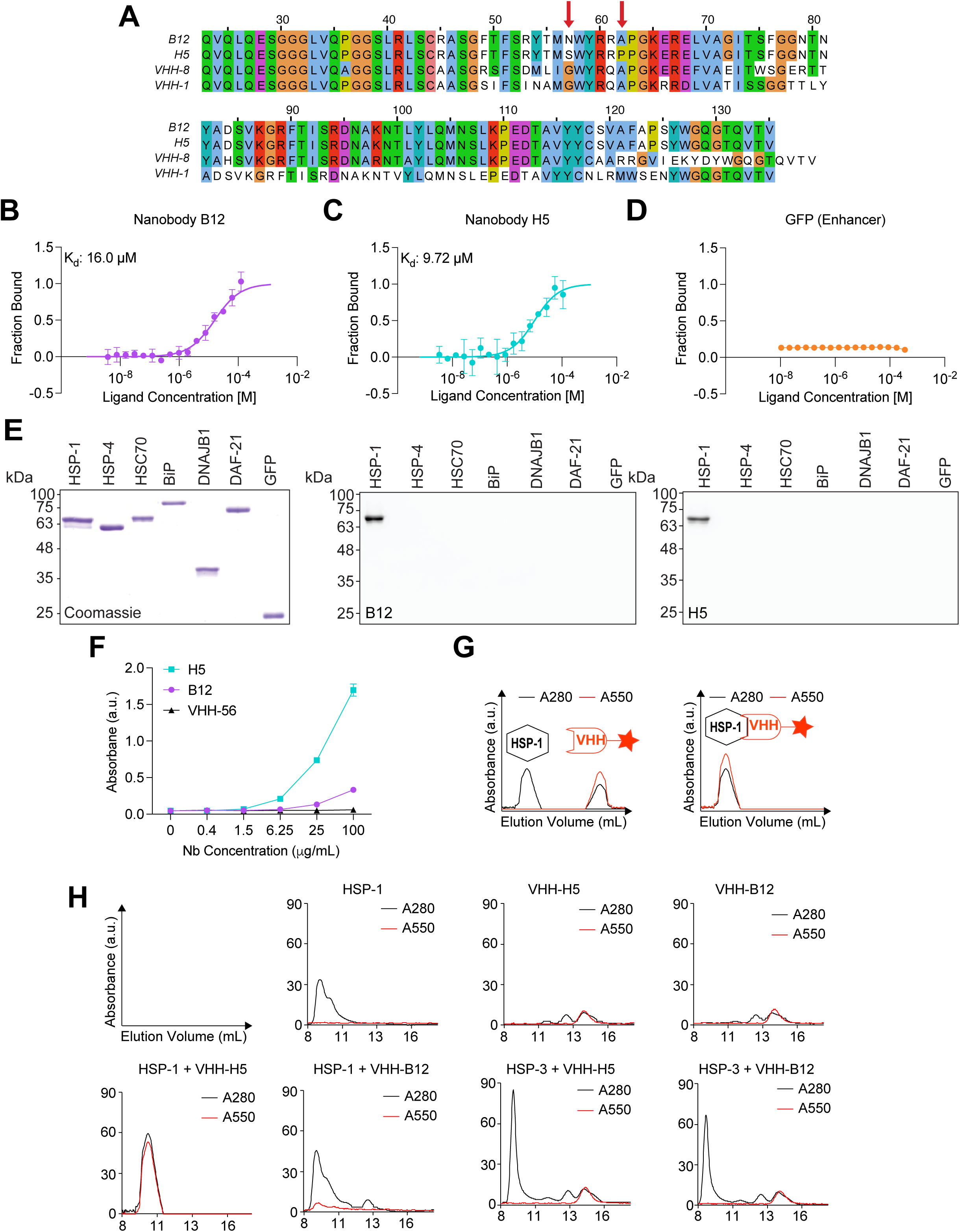
Generation of HSP-1-specific nanobodies B12 and H5. (A). Sequence alignments of nanobodies B12, H5, and two other nanobodies (VHH-1 and VHH-8) isolated from the same immune library without HSP-1 binding activity. Red arrows indicate differences in two amino acids between B12 and H5. Blue: Amino acids with hydrophobic side chains; Green: Amino acids with polar uncharged side chains; Purple: Amino acids with negatively charged side chains; Red: Amino acids with positively charged side chains; Other colors: special cases (B-D). MST binding curves for HSP-1 (20 nM) titrated against B12 (B, 0.0038–125 µM), H5 (C, 0.0033–109.4 µM) and VHH Enhancer (D, 0.0106–350 µM). Data represent mean ± SD of two replicates. (E). Coomassie stained SDS-PAGE analysis (left) and western blots using B12 (middle) or H5 (right) detecting the depicted chaperones and GFP. (F). Example ELISA using HSP-1 as the bait protein. (G). Schematic of size exclusion chromatography absorbance readouts. (H). Size exclusion chromatography of indicated combination of protein(s) and nanobody. For western blots and ELISAs, nanobodies were detected using the Precision Protein StrepTactin-HRP Conjugate (Bio-Rad, Cat #161038).

After cloning these sequences into an *E. coli* over-expression vector suitable for periplasmic nanobody expression (pHEN), we purified B12 and H5 and evaluated the specificity and binding capacity of each nanobody for recombinant HSP-1. Binding assays determined a K_d_ of 16.0 µM for B12 to HSP-1 and a K_d_ of 9.72 µM for H5 to HSP-1 (**Figs. 1B-C**). Importantly, control nanobody (anti-GFP; VHH_Enhancer_)^54^ did not bind to HSP-1 (**Fig. 1D**). Next, we determined the ability and specificity of biotinylated B12 and H5 to detect HSP-1 in western blot and ELISA assays. Both nanobodies recognized HSP-1 in these assays, confirming antigen targeting *in vitro* (**Figs. 1E-F, Supplemental Fig. 1A-D)**). Notably, neither B12 nor H5 detected other *C. elegans* or human HSP70 chaperones (e.g., HSP-4, HSC70, BiP), or any other non-HSP70 protein tested (e.g., DNAJB1, DAF-21/HSP90, GFP) in western blotting experiments (**Figs. 1E**). Notably, in ELISA assays, H5 demonstrated a noticeably stronger binding capability compared to B12 (**Fig. 1E, Supplemental Fig. 1D**), aligning with the enhanced binding affinity of H5 for properly folded HSP-1 suggested by the binding kinetics experiments (**Figs. 1B-C, F**)

To test B12 and H5 interactions with HSP-1 in an aqueous solution, we labelled both nanobodies with a C-terminal TAMRA fluorophore using sortase technology^55^ and analyzed nanobody-HSP-1 complex formation by analytic size exclusion chromatography (**Figs. 1G-H**). We observed stable binding of B12 and H5 to HSP-1 as indicated by the TAMRA signal eluting with the chaperone. Consistent with previous experiments, neither nanobody bound to HSP-3, a *C. elegans* BiP/HSPA5 ortholog (**Fig. 1H**).

These findings establish both B12 and H5 as selective binders of HSP-1, with H5 exhibiting stronger binding *in vitro*.

### B12 and H5 inhibit the ATPase and protein refolding capability of HSP-1

After establishing B12 and H5 bind to HSP-1 *in vitro*, we next wondered if these nanobodies could be used to modulate HSP-1 function. First, we sought to determine if B12 or H5 impacted the ATPase activity of HSP-1. *In vitro* ATPase assays showed that both B12 and H5 inhibited the DNJ-13-induced ATPase activity of HSP-1 in a dose-dependent manner. We observed significant reductions upon addition of 30 µM and a ∼50% reduction upon addition of 150 µM of either nanobody (**Figs. 2A-B**). Addition of an anti-GFP nanobody did not affect ATP hydrolysis or free phosphate production, again suggesting this result is specific to B12 and H5 (**Figs. 2A-B**).

**Figure 2.**
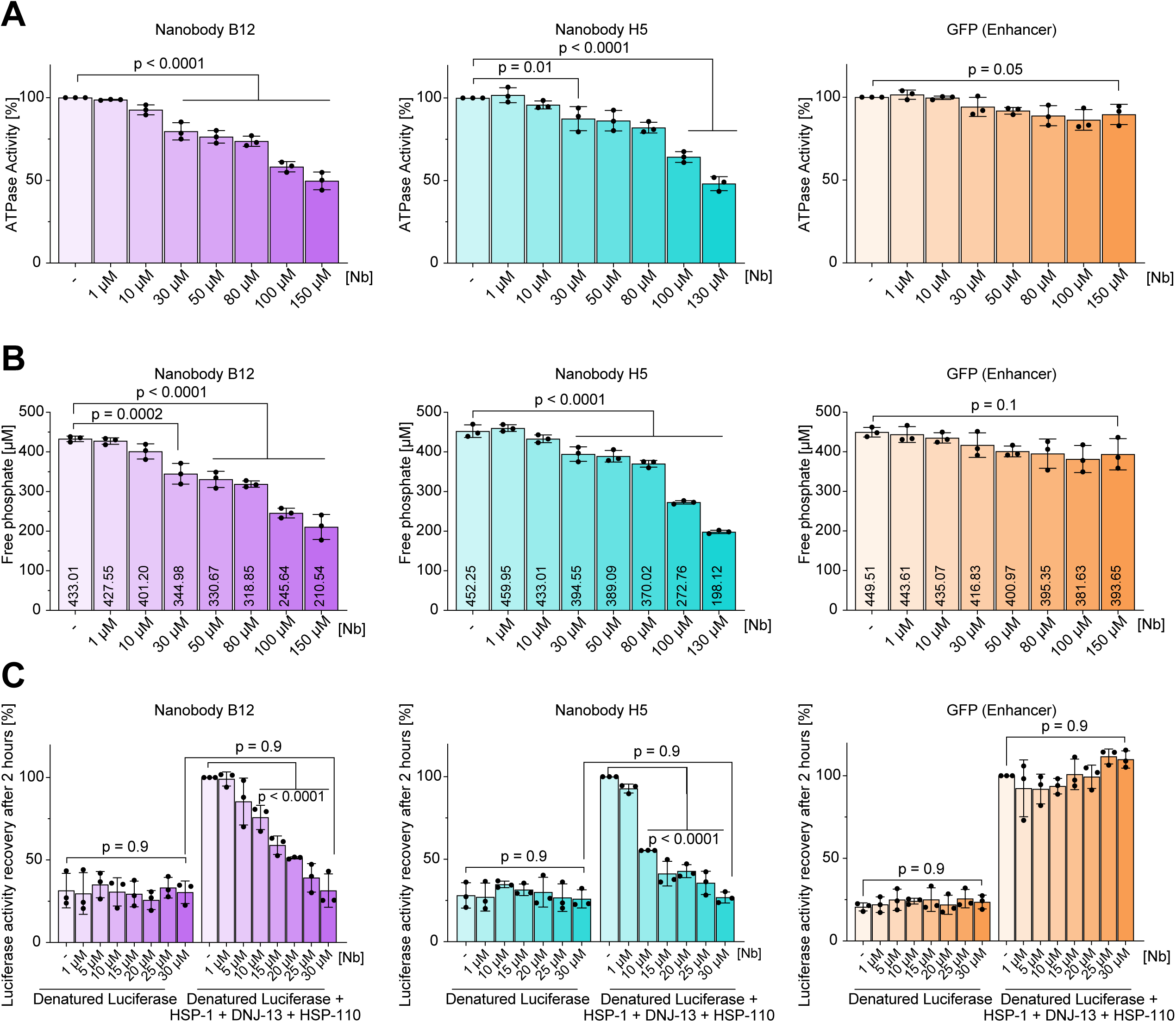
Nanobodies B12 and H5 inhibit HSP-1 function *in vitro*. (A) HSP-1-specific (A) ATPase assays and (B) quantifications of free-phosphate formation of reaction mixtures supplemented with indicated amount of nanobody B12, H5, or VHH Enhancer. Bars represent mean value of three replicates and error bars corresponds to the mean standard deviation. (C). HSP-1-driven Luciferase refolding assay supplemented with indicated amount of nanobody B12, H5, or VHH Enhancer. Luciferase activity was measured following a two-hour recovery and normalized to the values measured for the Trimeric-chaperone complex sample without nanobody. *Ordinary one-way ANOVA. p < 0.05 is considered statistically significant*.

Next, we assessed the ability of the nanobodies to affect refolding of denatured luciferase by HSP-1 and co-chaperones (DNJ-13, HSP-110)^26^. Interestingly, the addition of low micromolar (10-30 µM) concentrations of either B12 or H5 into this reaction limited luciferase refolding in a dose-dependent manner, with complete inhibition between 20-30 µM (**Fig. 2C**). The concentration of the nanobodies required to inhibit HSP-1 function aligned well with the obtained binding constants (**Fig. 1B-C**). However, the addition of the same concentrations of an anti-GFP (Enhancer) nanobody to the reaction mixture had no effect on luciferase refolding, suggesting this inhibition of HSP-1 refolding capability is specific to B12 and H5 (**Fig. 2C**).

Altogether, these data suggest B12 and H5 inhibit HSP-1 protein refolding capability and ATPase function *in vitro*.

### B12 and H5 bind to HSP-1 in complex *C. elegans* lysates

Antibodies and nanobodies are most useful if they recognize target antigens in complex samples, including cell or tissue lysates. We thus sought to determine if B12 and/or H5 nanobodies recognize endogenously expressed HSP-1. Using B12 or H5 as the primary antigen-binding moiety to detect HSP-1 in lysates via western blot, we found these nanobodies detected a single dominant band at the approximate molecular weight of HSP-1 (∼73 kDa) in wild-type *C. elegans* lysate but not in lysates of worms in which HSP-1 levels were depleted using *hsp-1* RNAi (**Fig. 3A-B, Supplemental Fig. 2A-C**). We also observed an *hsp-1* siRNA-sensitive faint band at ∼70 kDa migrating slightly faster than the dominant band, suggesting that B12 and H5 detect distinct HSP-1 populations that likely vary in their post-translational modification patterns (**Fig. 3A-B, Supplemental Fig. 2A-C**). In subsequent immunoprecipitation (IP) assays, we confirmed that biotinylated H5 and, to a lesser extent, B12, captured HSP-1 from the soluble fraction of total *C. elegans* lysates (**Fig. 3C**). These results confirm B12 and H5 detect endogenous HSP-1 in complex environments.

**Figure 3.**
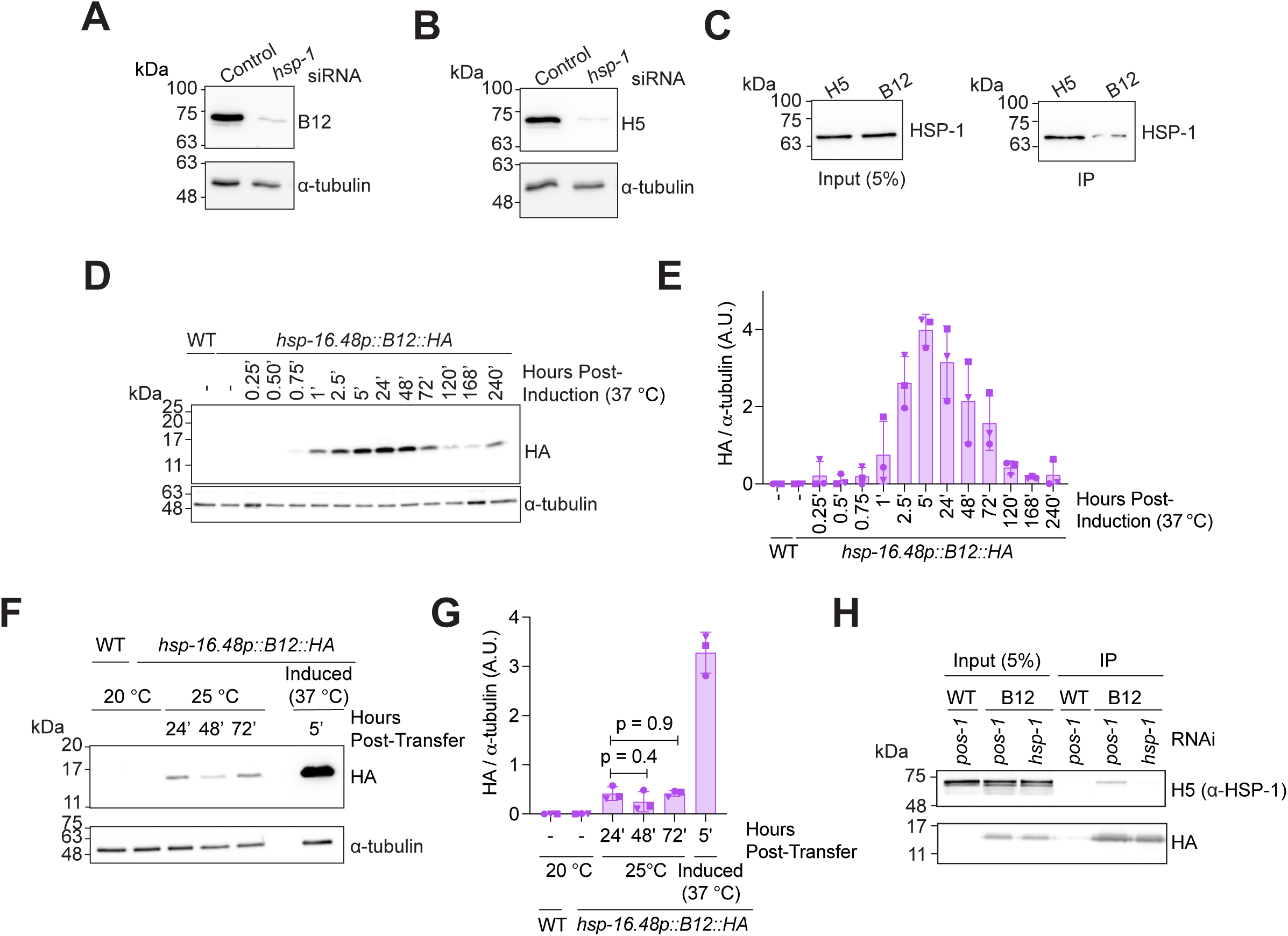
B12 and H5 detect HSP-1 from *C. elegans’* lysate. (A-B). Western blots of day 2 *C. elegans* that were grown on control (*pos-1* RNAi) plates and then transferred to either *pos-1* or *hsp-1* RNAi plates for 24 hours as day 1 adults. *pos-1* encodes a zinc-finger transcription factor required for embryonic development and sterilizes worms without affecting hatched animals^47–52^. Membranes were probed with either biotinylated (A) B12 or (B) H5 and α-tubulin (DSHB, Cat #12G10). (C). Western blots from immunoprecipitation assay using recombinant H5 or B12 as the primary antigen-binding agent and probed with an anti-HSC70 antibody (Proteintech, Cat #10654-1-AP) and α-tubulin (DSHB, Cat #12G10). (D-E). Example (D) western blot and (E) quantification of nanobody B12 expression (MTX298 (*hsp-16.48p::B12::HA*)) induced by a 30-minute heat shock at 37 °C (HA, Cell Signaling (C29F4) and α-tubulin (DSHB, Cat #12G10)). (F-G). Example (F) western blot and (G) quantification of nanobody B12 expression (MTX298 (*hsp-16.48p::B12::HA*)) placed at 25 °C for indicated amount of time or induced by 30-minute heat shock at 37 °C (HA, Cell Signaling (C29F4) and α-tubulin (DSHB, Cat #12G10). Induction was started in day 1 adults. (H). Co-immunoprecipitation assay of WT and MTX298 (“B12”) grown on indicated siRNA and collected 5 hours post 30-minute heat shock at 37 °C using magnetic anti-HA beads. *Ordinary one-way ANOVA. p < 0.05 is considered statistically significant*.

### Inducible expression of HSP-1-specific nanobodies is tolerated and bind to HSP-1 *in vivo*

We next generated transgenic *C. elegans* strains expressing nanobodies B12, H5, and VHH-7 containing a C-terminal HA-tag under control of the heat-inducible *hsp-16.48* promoter. VHH-7, a nanobody specific for Major Histocompatibility Complex class II^56^, served as non-targeting nanobody control. A 30-minute heat shock at 37 °C induced B12 expression, which peaked 5 hours post-induction and remained detectable for at least ten days (**Fig. 3D-E, Supplemental Fig. 3A**), whereas continuous cultivation of worms at 25 °C, a condition known to induce modest heat stress, generated basal expression at approximately 16 % of peak level (**Fig. 3F-G, Supplemental Fig. 3B**). We observed comparable expression patterns for nanobodies H5 and VHH-7, although the magnitude of expression varied depending on the specific nanobody-expressing strain and was strongest in the B12-expressing strain (**Supplemental Fig. 3C-E**). These results demonstrate that camelid nanobodies can be efficiently expressed in transgenic *C. elegans*. Using magnetic agarose beads conjugated with anti-HA antibodies, we also showed that pulling on the HA-tag of B12 allowed for the co-immunoprecipitation of the B12-HSP-1 complex from worm lysates (**Fig. 3H, Supplemental Fig. 3F**). Altogether, these data indicate that *in vivo* expressed B12 nanobodies interact with HSP-1.

### *In vivo* expression of B12, but not H5, reduces survival and proteotoxic stress resistance in *C. elegans*

Given that B12 and H5 inhibit HSP-1 chaperone function *in vitro* (**Fig. 2**) and that they recognize endogenous HSP-1 (**Fig. 3**), we sought to test if these nanobodies modulate HSP-1 activity in living *C. elegans*. To express and sustain nanobody expression without inducing a severe heat stress, we transferred adult WT and nanobody-expressing animals to 25 °C and measured their survival. Low-level B12 expression at 25 °C significantly shortened adult survival, matching the reduction in survival observed when *hsp-1* was knocked down in WT animals using RNAi (**Fig. 4A, Supplemental Fig. 4A**). In contrast, H5 or control-nanobody expressing animals kept at 25 °C did not exhibit a decrease in survival compared to WT (**Figs. 4B-C, Supplemental Fig. 4B-C**). Survival was unchanged in B12 or H5-expressing animals following a 30-minute induction at 37 °C compared to WT animals (**Supplemental Figs. 4D-E**).

**Figure 4.**
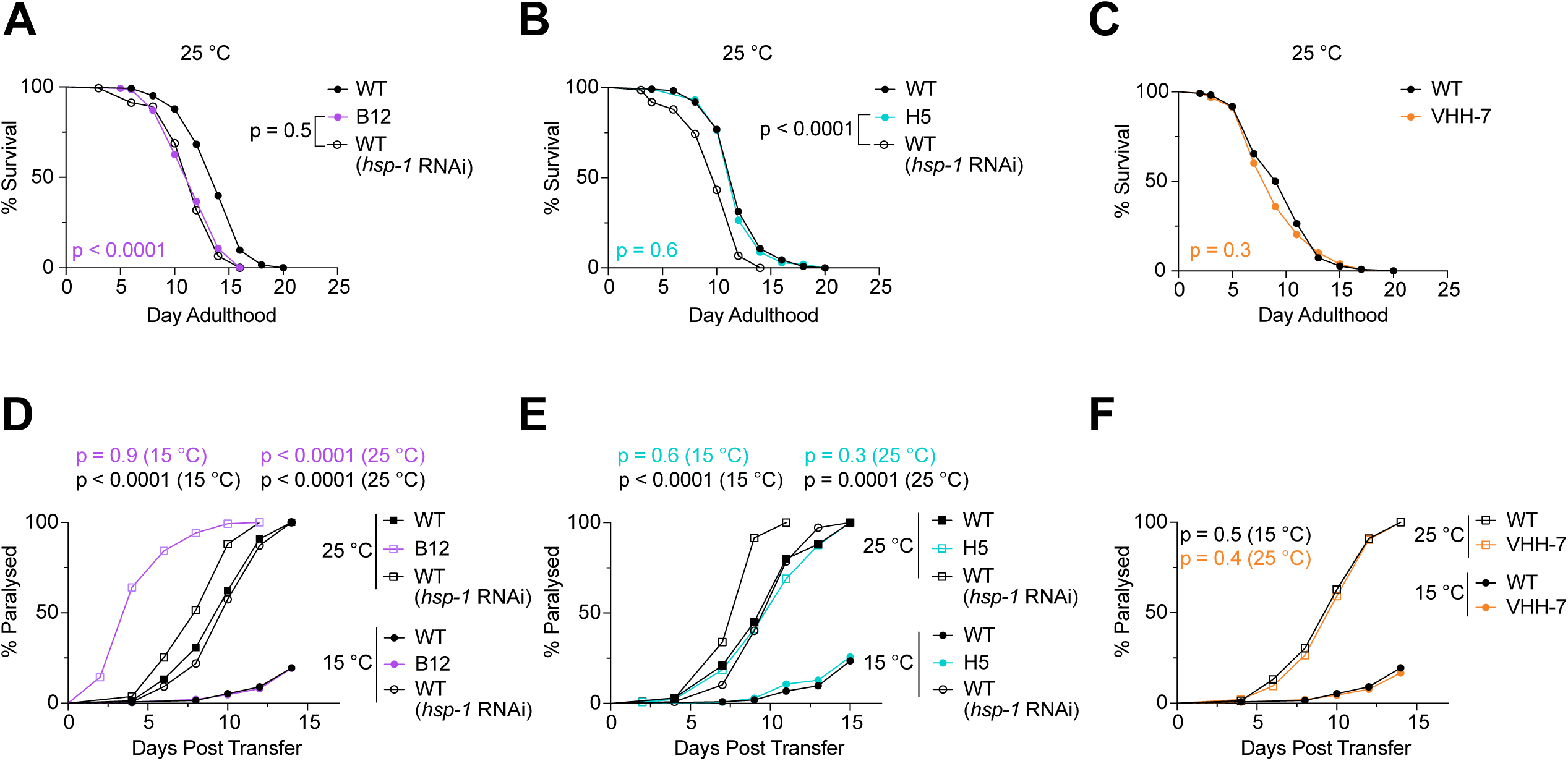
Mild expression of B12 mimics phenotypes of *hsp-1* knockdown in *C. elegans*. (A-C). Survival curves of animals placed at 25 °C beginning on day 1 of adulthood. (D-F). Paralysis curves of Aβ_1–42_-expressing animals at indicated temperature. For A-C, WT: wild type; B12: MTX298 (*hsp-16.48p::B12::HA*); H5: MTX277 (*hsp-16.48p::H5::HA*); VHH-7: MT24421 (*hsp-16.48p:: VHH-7::HA*). For D-F, WT: GMC101 (*dvIs100*); B12: MTX307 (*mtmIs20; dvIs100*),); H5: MTX319 (*mtmEx103; dvIs100*),; MTX329 (*nIs775; dvIs100*). Number of animals, median lifespan, and median day of paralysis are shown in Supplemental Tables 2 and 3. Replicate experiments are shown in Supplemental Figures 4 and 5. *Log-rank Mantel-Cox test. p < 0.05 is considered statistically significant*.

We then wondered if nanobody expression altered the worm’s ability to mitigate protein misfolding stress. To test this prediction, we introduced the nanobody-encoding alleles into a strain expressing human Aβ_1–42_ in body-wall muscle cells (GMC101 (*dvIs100 [unc-54p:: Aβ_1–42_::unc-54 3’-UTR + mtl-2p::GFP])*^57^). When placed at the inducive temperature of 25 °C, these animals progressively lose motor function due to the accumulation of misfolded Aβ ^57^. We hypothesized that B12 and H5 expression would accelerate paralysis in a GMC101 background. Indeed, at 25 °C we observed B12-expressing animals paralyzed significantly faster than wild type worms on both control and *hsp-1* siRNA at the same temperature (**Fig. 4D, Supplemental Fig. 5A**). Interestingly, there was no difference in paralysis between B12 and WT animals at the non-inducive temperature (15 °C), in which the *hsp-16.48* promoter is almost entirely inactive; this was unlike WT animals at the non-inducive temperature on *hsp-1* siRNA, which showed significantly increased rates of paralysis (**Fig. 4D, Supplemental Fig. 5A**). Notably, no comparable differences in paralysis were observed at both the non-inducive and inducive temperatures in animals which expressed H5 or VHH-7 (**Fig. 4E-F Supplemental Fig. 5B-C**).

Collectively, these results demonstrate low-level expression of B12 effectively suppresses HSP-1 activity *in vivo* and phenocopies *hsp-1* knockdown. Our results further confirm the necessity of HSP-1 for mild heat stress survival and proteotoxic stress-resistance in *C. elegans*.

## Discussion

In this study, we identified and characterized two novel HSP-1-specific nanobodies (B12 and H5). We show that these nanobodies, which differ by only two amino acids in their CDR1 region, bind to HSP-1 both *in vitro* and *in vivo* (**Figs. 1, 3**). Using HSP-1-specific ATPase and luciferase refolding assays, we show that B12 and H5 inhibit the ATPase activity, as well as the ability of HSP-1 to refold misfolded protein (luciferase), in a dose-dependent manner (**Fig. 2**). Finally, we show that low level *in vivo* expression of B12 (**Fig. 3**) shortens survival and reduces proteotoxic stress resistance in *C. elegans* —replicating the phenotypes of *hsp-1* knockdown (**Fig. 4**). Overall, these results demonstrate the effectiveness and specificity of these nanobodies to inhibit a specific HSP70 family chaperone *in vitro* and *in vivo*.

The ability to modulate HSP70 chaperone levels or activity is essential to understanding the physiological functions of these proteins in aging and protein-misfolding diseases. By precisely controlling HSP70 function in living organisms, we can begin to delineate the specific roles of HSP70 chaperones in disease processes and potentially develop novel targeted therapeutics. However, the ability to modulate *in vivo* chaperone activity remains difficult, since many HSP70s are essential and thus hard to target using conventional genetic approaches. Furthermore, the conserved structural similarity^58^ of HSP70 chaperones makes finding protein-specific small molecule modulators difficult to utilize. Thus, the temporal expression of nanobodies which effectively modulate the function of a specific HSP70 chaperone presents an exciting and effective strategy to advance our knowledge of the functions of specific chaperones in different physiological and pathophysiological contexts.

Interestingly, we observed that, *in* vivo, B12 is more potent to reduce HSP-1 activity than H5. This discrepancy may at least in part be explained by differences in absolute nanobody levels upon transgene induction. Despite utilizing the same promoter to express B12, H5, and VHH-7 in *C. elegans*, transgene copy number, chromosomal position effects, partial epigenetic silencing, integrated versus extrachromosomal arrays, and differences in mRNA stability or translation efficiency likely prevent to achieve uniform transgene expression levels across strains^59^.

Overall, our data demonstrate the effectiveness of using nanobodies to modulate HSP70 family chaperone activity *in vitro* and *in vivo*.

## Study Limitations

Our approach of expressing B12 and H5 under a stress-induced promoter is effective as it provides robust protein expression. However, it does limit the experimental design to assays which maintain nanobody expression under what could be considered “stressful” conditions (e.g., elevated temperature); thus, interpretations as to the basal physiological roles of HSP-1 may be complex to interpret. In the future, this may be circumvented using other inducible technologies, such as tetracycline-inducible systems (e.g., Tet/Q hybrid system)^60^, optogenetic systems^61^, or chemically induced systems^62,63^, which would allow for temporal expression without the need for a canonical mild stressor. However, these require the assembly of more complex transgenics and/or specialized equipment, thus limiting their ease of generation and use. Regardless, our results demonstrating B12 expression increases misfolding-induced paralysis at 25 °C but has no effect at the non-inducive 15 °C (**Fig. 4D**) clearly shows the effectiveness of this current system while providing a base for future tool generation and expansion.

## Acknowledgments

We thank the members of the Truttmann and Kirstein labs for helpful comments and discussion. William Giblin is acknowledged for proof-reading the manuscript draft and integrating nanobody-expressing strains. NDU was supported by Training Grants GM008322-30 and AG000114-37, as well as award 1F31AG085891-01A1. MCT is supported by an Alzheimer’s foundation Young Investigator Award, a Ruth K Broad foundation award and grant 1R35GM142561.

## Author contributions

MCT, JK, and NDU supervised the project. NDU and MCT designed and planned the experiments. NDU, KG, ZM, AS, AK, and ST performed all experiments. NDU wrote the initial draft of the manuscript, which was subsequently edited by NDU, JK and MCT.

## Data availability statement

The unprocessed raw datasets generated and analyzed during the current study are available from the corresponding author upon reasonable request.

## Additional information

The authors declare no competing interests.

---

**Supplemental Figure S1.**
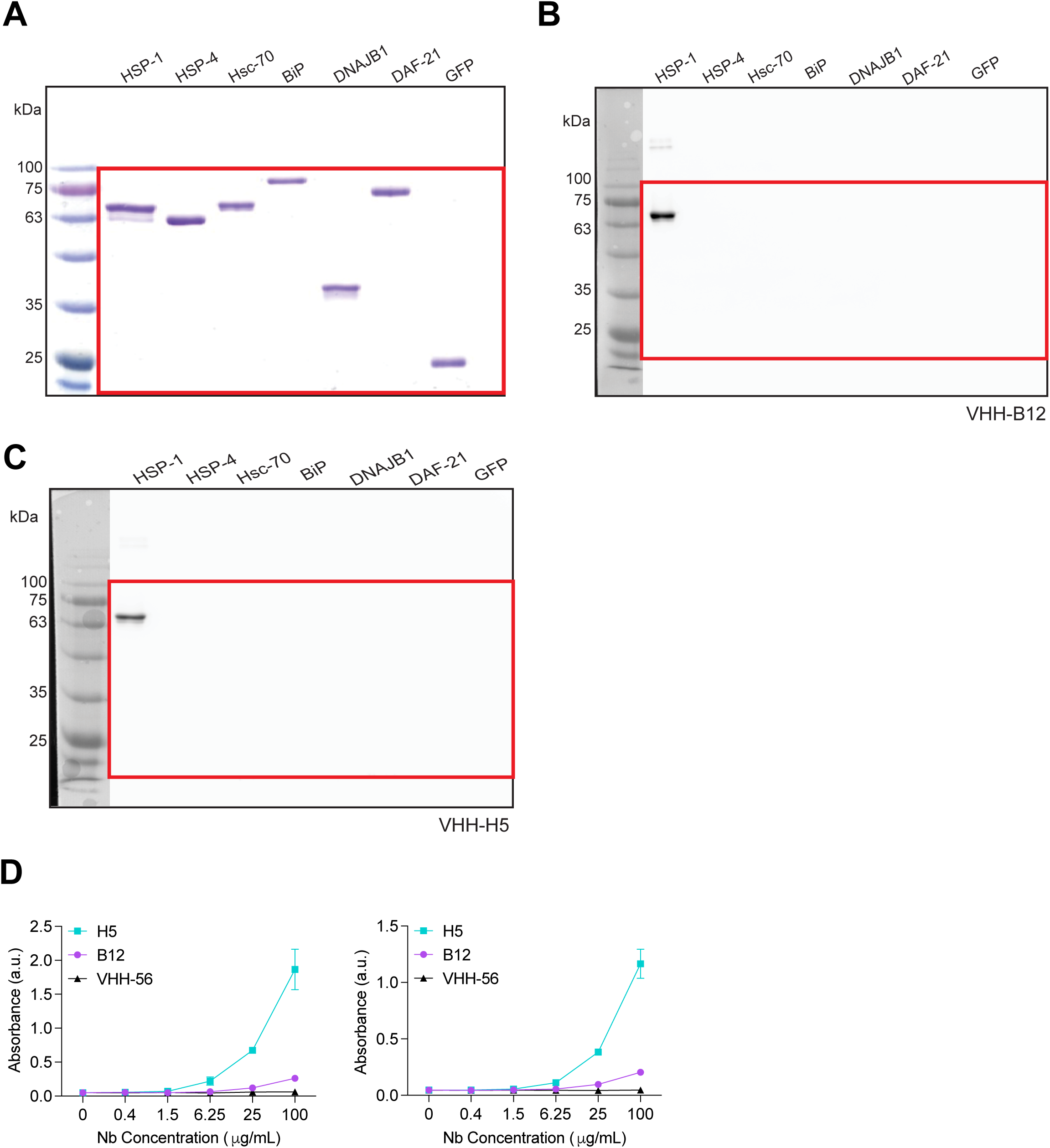
B12 and H5 recognize HSP-1 in Western blots and ELISA assays. (A-C) Uncropped Coomassie stain and western blot example of blots shown in Figure 1. (D). Two additional replicates of ELISA assays using HSP-1 as the bait protein. Nanobodies were conjugated with biotin using sortase technology and detected using the Precision Protein StrepTactin-HRP Conjugate (BioRad, Cat #161038). Red box indicates what is shown in main text.

**Supplemental Figure S2.**
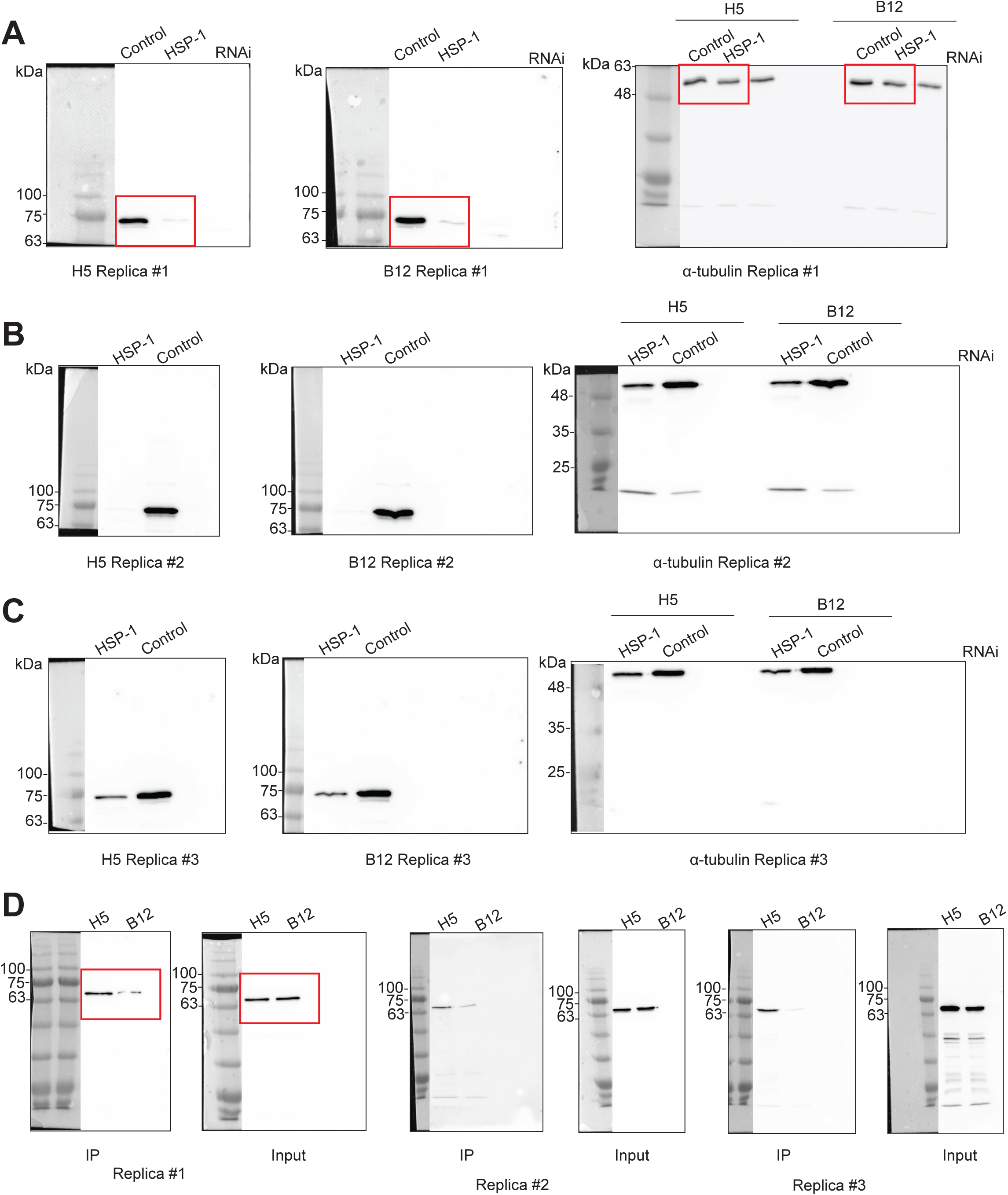
B12 and H5 recognize endogenous HSP-1. (A-C). Three biological replicates of western blots showing biotinylated B12 and H5 detect HSP-1 in worm lysate. (D). Three biological replicates of western blots from immunoprecipitation assays using recombinant H5 or B12 as the primary antigen-binding agent. Membranes were probed with an anti-HSC70 antibody (Proteintech, Cat #10654-1-AP) and α-tubulin (DSHB, Cat #12G10). Red box indicates what is shown in Figure 3.

**Supplemental Figure S3.**
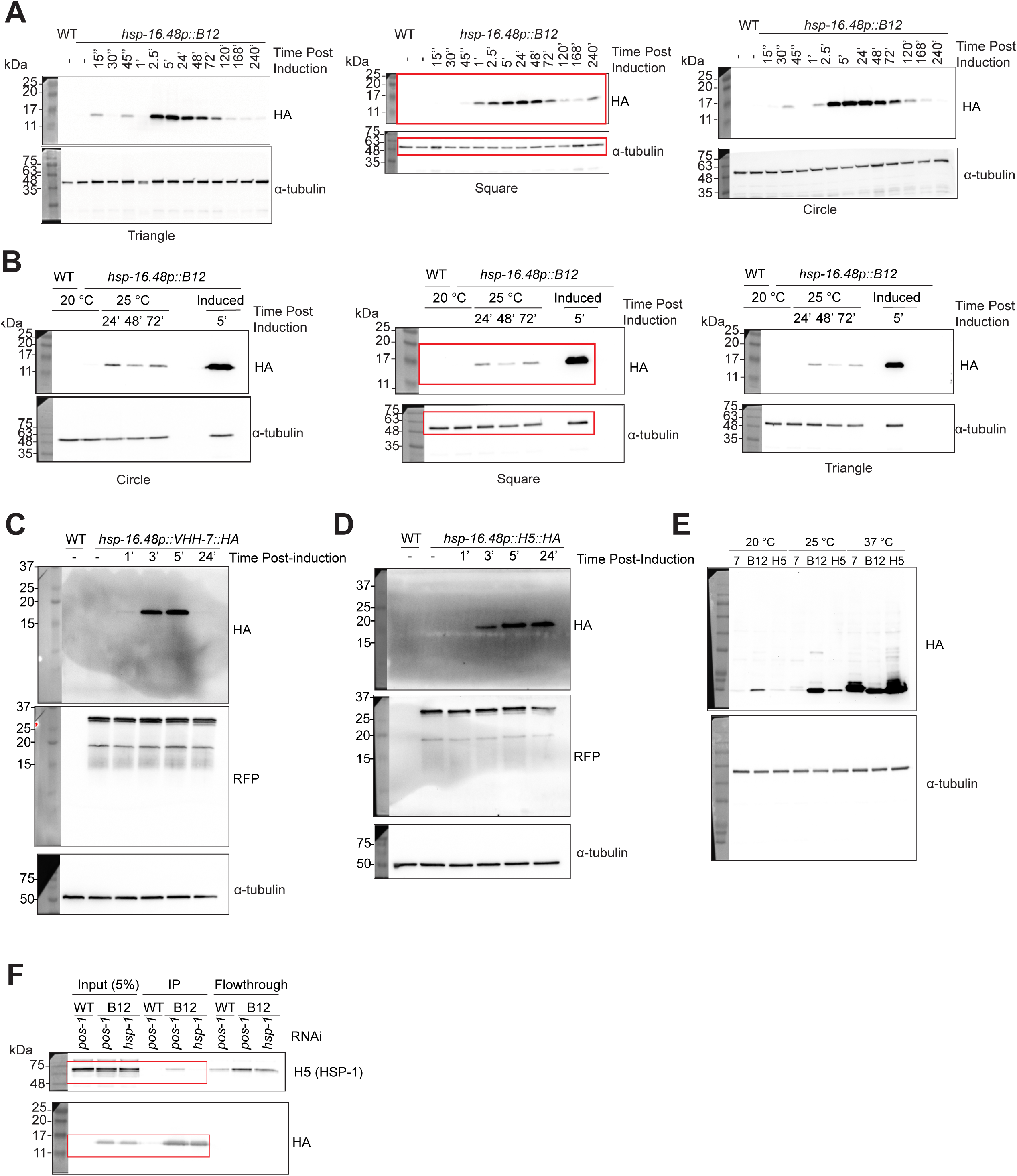
HSP-1-specific nanobodies can be expressed endogenously in *C. elegans*. (A). Western blot replicates of time course assay for MTX298 (*hsp-16.48p::B12::HA*). Protein induction was induced by a 30-minute heat shock at 37 °C. (B). Western blot replicates of time course assay of MTX298 (*hsp-16.48p::B12::HA*) protein collected after placing day 1 adult animals at 25 °C for indicated amount of time or 5 hours after a 30-minute heat shock at 37 °C (“Induced”). (C-D). Western blots for protein expression time course assays of (C) MTX277 (*hsp-16.48p::H5::HA*) and (D) MT24421 (*hsp-16.48p:: VHH-7::HA*). Nanobody induction was driven by a 30-minute heat shock at 37 °C. (E). Western blot comparing induction of MTX298 (B12), MTX277 (H5) and MT24421 (7) at indicated temperatures. Animals were synchronized and eggs were placed directly at either 20 °C or 25 °C and collected 72 hours later or induced as day 1 adults by a 30-minute heat shock at 37 °C and collected after 5 hours. (F). Uncropped immunoprecipitation assay of day 2 adult worms collected 5 hours after induction. Day 1 adults were transferred to either *pos-1* or *hsp-1* siRNA 24 hours prior to induction. “B12” is MTX298. HA, Cell Signaling (Cat #C29F4); RFP Proteintch (Cat #6g6), α-tubulin (DSHB, Cat #12G10). Red box indicates what is shown in Figure 3.

**Supplemental Figure S4.**
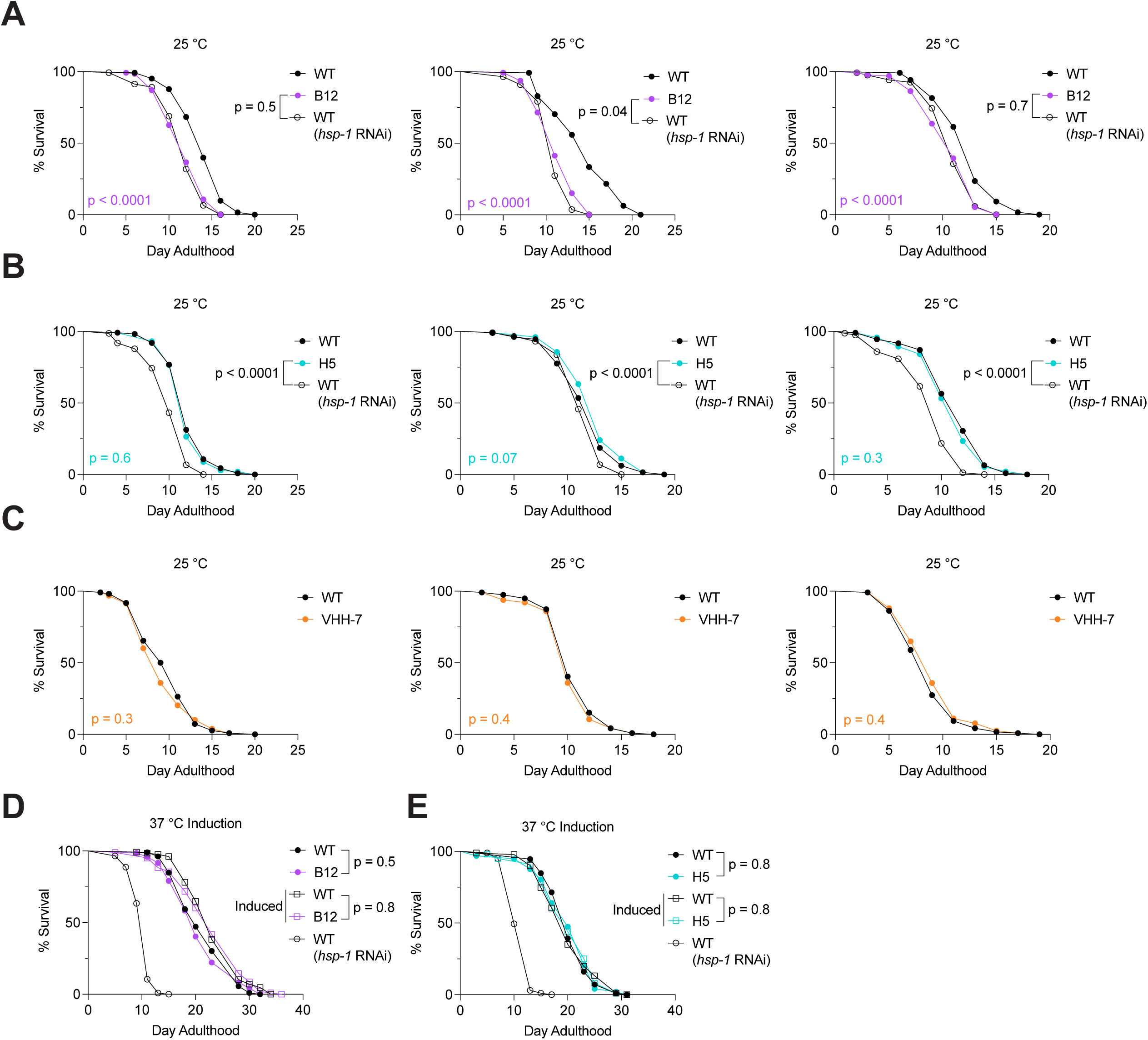
*C. elegans* lifespan assays. (A-C). Replicates of survival curves for (A) MTX298, (B) MTX277, and (C) MT24421 which were grown for 72 hours at 20 °C then transferred to 25 °C as day 1 adults. (D-E). Survival curves of (D) MTX298 and (E) MTX277 following a 30-minute induction at 37 °C. Day 1 adult animals were induced. For all experiments WT animals were transferred to *hsp-1* siRNA as day 1 adults. WT: wild type; B12: MTX298 (*hsp-16.48p::B12::HA*); H5: MTX277 (*hsp-16.48p::H5::HA*); VHH-7: MT24421 (*hsp-16.48p:: VHH-7::HA*). *Log-rank Mantel-Cox test. p < 0.05 is considered statistically significant*. Median lifespan and number of animals for all experiments are listed in Supplemental Table 2.

**Supplemental Figure S5.**
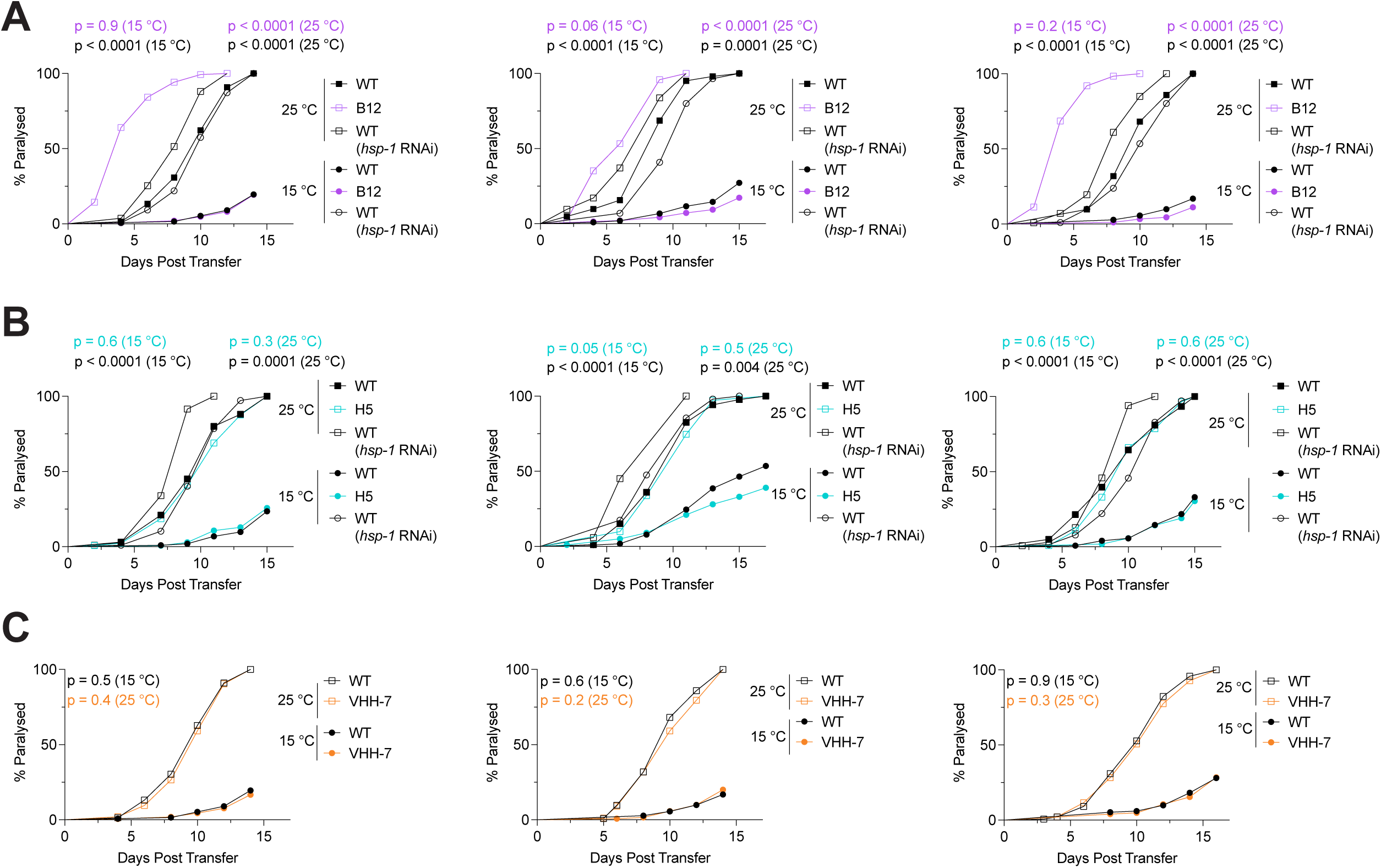
*C. elegans* paralysis assays. (A-C). Replicates of paralysis assays for (A) MTX307 (“B12”, *hsp-16.48p::B12::HA; dvIs100*), (B) MTX320 (“H5”, *hsp-16.48p::H5::HA; dvIs100*), and (C) MTX329 (“VHH-7”, *hsp-16.48p::VHH-7::HA; dvIs100*) which were grown for 72 hours at 20 °C then transferred to 25 °C as day 1 adults. WT: GMC101 (*dvIs100 [unc-54p::A-beta-1-42::unc-54 3’-UTR + mtl-2p::GFP]*). *Log-rank Mantel-Cox test. p < 0.05 is considered statistically significant*. Median paralysis and number of animals per experiment is listed in Supplemental Table 3.

## References

1. Yu E meng, Yoshinaga T, Jalufka FL, Ehsan H, Mark Welch DB, Kaneko G. The complex evolution of the metazoan HSP70 gene family. Sci Rep. 2021;11(1):17794. doi:10.1038/s41598-021-97192-9

2. Flaherty KM, DeLuca-Flaherty C, McKay DB. Three-dimensional structure of the ATPase fragment of a 70K heat-shock cognate protein. Nature. 1990;346(6285):623-628. doi:10.1038/346623a0

3. Zhu X, Zhao X, Burkholder WF, et al. Structural Analysis of Substrate Binding by the Molecular Chaperone DnaK. Science. 1996;272(5268):1606-1614. doi:10.1126/science.272.5268.1606

4. Swain JF, Schulz EG, Gierasch LM. Direct Comparison of a Stable Isolated Hsp70 Substrate-binding Domain in the Empty and Substrate-bound States. Journal of Biological Chemistry. 2006;281(3):1605–1611. doi:10.1074/jbc.M509356200

5. Oshiro N, Takahashi R, Yoshino K ichi, et al. The Proline-rich Akt Substrate of 40 kDa (PRAS40) Is a Physiological Substrate of Mammalian Target of Rapamycin Complex 1. Journal of Biological Chemistry. 2007;282(28):20329–20339. doi:10.1074/jbc.M702636200

6. Mayer MP, Bukau B. Hsp70 chaperones: cellular functions and molecular mechanism. Cell Mol Life Sci. 2005;62(6):670–684. doi:10.1007/s00018-004-4464-6

7. Rohland L, Kityk R, Smalinskaitė L, Mayer MP. Conformational dynamics of the Hsp70 chaperone throughout key steps of its ATPase cycle. Proc Natl Acad Sci U S A. 2022;119(48):e2123238119. doi:10.1073/pnas.2123238119

8. Kampinga HH, Craig EA. The HSP70 chaperone machinery: J proteins as drivers of functional specificity. Nat Rev Mol Cell Biol. 2010;11(8):579–592. doi:10.1038/nrm2941

9. Misselwitz B, Staeck O, Rapoport TA. J Proteins Catalytically Activate Hsp70 Molecules to Trap a Wide Range of Peptide Sequences. Molecular Cell. 1998;2(5):593–603. doi:10.1016/S1097-2765(00)80158-6

10. Karzai AW, McMacken R. A bipartite signaling mechanism involved in DnaJ-mediated activation of the Escherichia coli DnaK protein. J Biol Chem. 1996;271(19):11236–11246. doi:10.1074/jbc.271.19.11236

11. Gupta A, Bansal A, Hashimoto-Torii K. HSP70 and HSP90 in neurodegenerative diseases. Neurosci Lett. 2020;716:134678. doi:10.1016/j.neulet.2019.134678

12. Witt SN. Hsp70 molecular chaperones and Parkinson’s disease. Biopolymers. 2010;93(3):218–228. doi:10.1002/bip.21302

13. Turturici G, Sconzo G, Geraci F. Hsp70 and its molecular role in nervous system diseases. Biochem Res Int. 2011;2011:618127. doi:10.1155/2011/618127

14. Murphy ME. The HSP70 family and cancer. Carcinogenesis. 2013;34(6):1181–1188. doi:10.1093/carcin/bgt111

15. Abe M, Manola JB, Oh WK, et al. Plasma Levels of Heat Shock Protein 70 in Patients with Prostate Cancer: A Potential Biomarker for Prostate Cancer. Clinical Prostate Cancer. 2004;3(1):49–53. doi:10.3816/CGC.2004.n.013

16. Bobkova NV, Garbuz DG, Nesterova I, et al. Therapeutic Effect of Exogenous Hsp70 in Mouse Models of Alzheimer’s Disease. JAD. 2013;38(2):425–435. doi:10.3233/JAD-130779

17. Radons J. The human HSP70 family of chaperones: where do we stand? Cell Stress and Chaperones. 2016;21(3):379–404. doi:10.1007/s12192-016-0676-6

18. Hartl FU, Hayer-Hartl M. Molecular Chaperones in the Cytosol: from Nascent Chain to Folded Protein. Science. 2002;295(5561):1852–1858. doi:10.1126/science.1068408

19. Brocchieri L, Conway De Macario E, Macario AJ. hsp70 genes in the human genome: Conservation and differentiation patterns predict a wide array of overlapping and specialized functions. BMC Evol Biol. 2008;8(1):19. doi:10.1186/1471-2148-8-19

20. Liu T, Daniels CK, Cao S. Comprehensive review on the HSC70 functions, interactions with related molecules and involvement in clinical diseases and therapeutic potential. Pharmacol Ther. 2012;136(3):354–374. doi:10.1016/j.pharmthera.2012.08.014

21. Dastoor Z, Dreyer JL. Nuclear translocation and aggregate formation of heat shock cognate protein 70 (Hsc70) in oxidative stress and apoptosis. Journal of Cell Science. 2000;113(16):2845–2854. doi:10.1242/jcs.113.16.2845

22. Tsukahara F, Maru Y. Identification of novel nuclear export and nuclear localization-related signals in human heat shock cognate protein 70. J Biol Chem. 2004;279(10):8867–8872. doi:10.1074/jbc.M308848200

23. Bański P, Mahboubi H, Kodiha M, Shrivastava S, Kanagaratham C, Stochaj U. Nucleolar targeting of the chaperone hsc70 is regulated by stress, cell signaling, and a composite targeting signal which is controlled by autoinhibition. J Biol Chem. 2010;285(28):21858–21867. doi:10.1074/jbc.M110.117291

24. Wang F, Bonam SR, Schall N, et al. Blocking nuclear export of HSPA8 after heat shock stress severely alters cell survival. Sci Rep. 2018;8(1):16820. doi:10.1038/s41598-018-34887-6

25. Truttmann MC, Pincus D, Ploegh HL. Chaperone AMPylation modulates aggregation and toxicity of neurodegenerative disease-associated polypeptides. Proc Natl Acad Sci USA. 2018;115(22). doi:10.1073/pnas.1801989115

26. Kirstein J, Arnsburg K, Scior A, et al. In vivo properties of the disaggregase function of J-proteins and Hsc70 in Caenorhabditis elegans stress and aging. Aging Cell. 2017;16(6):1414–1424. doi:10.1111/acel.12686

27. Ciechanover A, Kwon YT. Protein Quality Control by Molecular Chaperones in Neurodegeneration. Front Neurosci. 2017;11:185. doi:10.3389/fnins.2017.00185

28. Kanack AJ, Olp MD, Newsom OJ, et al. Chemical Regulation of the Protein Quality Control E3 Ubiquitin Ligase C-Terminus of Hsc70 Interacting Protein (CHIP). Chembiochem. 2022;23(6):e202100633. doi:10.1002/cbic.202100633

29. Abildgaard AB, Voutsinos V, Petersen SD, et al. HSP70-binding motifs function as protein quality control degrons. Cell Mol Life Sci. 2023;80(1):32. doi:10.1007/s00018-022-04679-3

30. Rebeaud ME, Mallik S, Goloubinoff P, Tawfik DS. On the evolution of chaperones and cochaperones and the expansion of proteomes across the Tree of Life. Proc Natl Acad Sci USA. 2021;118(21):e2020885118. doi:10.1073/pnas.2020885118

31. Hamers-Casterman C, Atarhouch T, Muyldermans S, et al. Naturally occurring antibodies devoid of light chains. Nature. 1993;363(6428):446-448. doi:10.1038/363446a0

32. Liu X, Wang Y, Sun L, et al. Screening and optimization of shark nanobodies against SARS-CoV-2 spike RBD. Antiviral Res. 2024;226:105898. doi:10.1016/j.antiviral.2024.105898

33. Spinelli S, Frenken L, Bourgeois D, et al. The crystal structure of a llama heavy chain variable domain. Nat Struct Mol Biol. 1996;3(9):752–757. doi:10.1038/nsb0996-752

34. Desmyter A, Transue TR, Ghahroudi MA, et al. Crystal structure of a camel single-domain VH antibody fragment in complex with lysozyme. Nat Struct Mol Biol. 1996;3(9):803–811. doi:10.1038/nsb0996-803

35. Chen WH, Hajduczki A, Martinez EJ, et al. Shark nanobodies with potent SARS-CoV-2 neutralizing activity and broad sarbecovirus reactivity. Nat Commun. 2023;14(1):580. doi:10.1038/s41467-023-36106-x

36. Zhu H, Ding Y. Nanobodies: From Discovery to AI-Driven Design. Biology (Basel). 2025;14(5):547. doi:10.3390/biology14050547

37. Arbabi Ghahroudi M, Desmyter A, Wyns L, Hamers R, Muyldermans S. Selection and identification of single domain antibody fragments from camel heavy-chain antibodies. FEBS Letters. 1997;414(3):521–526. doi:10.1016/S0014-5793(97)01062-4

38. Zhou X, Hao R, Chen C, et al. Rapid Delivery of Nanobodies/VHHs into Living Cells via Expressing In Vitro-Transcribed mRNA. Mol Ther Methods Clin Dev. 2020;17:401–408. doi:10.1016/j.omtm.2020.01.008

39. Zhang J, Sun H, Pei W, Jiang H, Chen J. Nanobody-based immunosensing methods for safeguarding public health. J Biomed Res. 2021;35(4):318–326. doi:10.7555/JBR.35.20210108

40. Cater JH, El Salamouni NS, Mansour GH, et al. Optimised nanobody-based quenchbodies for enhanced protein detection. Commun Biol. 2025;8(1):937. doi:10.1038/s42003-025-08359-3

41. Teunissen AJP, Abousaway OB, Munitz J, et al. Employing nanobodies for immune landscape profiling by PET imaging in mice. STAR Protoc. 2021;2(2):100434. doi:10.1016/j.xpro.2021.100434

42. Zhao S, Luo J, Xu P, et al. Designed peptide binders and nanobodies as PROTAC starting points for targeted degradation of PCNA and BCL6. International Journal of Biological Macromolecules. 2025;308:142667. doi:10.1016/j.ijbiomac.2025.142667

43. Truttmann MC, Wu Q, Stiegeler S, Duarte JN, Ingram J, Ploegh HL. HypE-specific nanobodies as tools to modulate HypE-mediated target AMPylation. J Biol Chem. 2015;290(14):9087–9100. doi:10.1074/jbc.M114.634287

44. Nillegoda NB, Kirstein J, Szlachcic A, et al. Crucial HSP70 co-chaperone complex unlocks metazoan protein disaggregation. Nature. 2015;524(7564):247-251. doi:10.1038/nature14884

45. Scior A, Buntru A, Arnsburg K, et al. Complete suppression of Htt fibrilization and disaggregation of Htt fibrils by a trimeric chaperone complex. The EMBO Journal. 2018;37(2):282–299. doi:10.15252/embj.201797212

46. Ayala Mariscal SM, Pigazzini ML, Richter Y, et al. Identification of a HTT-specific binding motif in DNAJB1 essential for suppression and disaggregation of HTT. Nat Commun. 2022;13(1):4692. doi:10.1038/s41467-022-32370-5

47. Urban ND, Lacy SM, Van Pelt KM, et al. Functionally diversified BiP orthologs control body growth, reproduction, stress resistance, aging, and ER-Phagy in *Caenorhabditis elegans*. Published online January 19, 2025. doi:10.1101/2025.01.14.633073

48. Tabara H, Hill RJ, Mello CC, Priess JR, Kohara Y. pos-1 encodes a cytoplasmic zinc-finger protein essential for germline specification in C. elegans. Development. 1999;126(1):1–11. doi:10.1242/dev.126.1.1

49. Tabara H, Sarkissian M, Kelly WG, et al. The rde-1 gene, RNA interference, and transposon silencing in C. elegans. Cell. 1999;99(2):123–132. doi:10.1016/s0092-8674(00)81644-x

50. Farley BM, Pagano JM, Ryder SP. RNA target specificity of the embryonic cell fate determinant POS-1. RNA. 2008;14(12):2685–2697. doi:10.1261/rna.1256708

51. Taylor MN, Spandana Boddu S, Vega NM. Using Single-Worm Data to Quantify Heterogeneity in Caenorhabditis elegans-Bacterial Interactions. J Vis Exp. 2022;(185). doi:10.3791/64027

52. Van Pelt KM, Truttmann MC. Loss of FIC-1-mediated AMPylation activates the UPR^ER^ and upregulates cytosolic HSP70 chaperones to suppress polyglutamine toxicity. Published online November 28, 2024. doi:10.1101/2024.11.27.625751

53. Rual JF, Ceron J, Koreth J, et al. Toward improving Caenorhabditis elegans phenome mapping with an ORFeome-based RNAi library. Genome Res. 2004;14(10B):2162–2168. doi:10.1101/gr.2505604

54. Kirchhofer A, Helma J, Schmidthals K, et al. Modulation of protein properties in living cells using nanobodies. Nat Struct Mol Biol. 2010;17(1):133–138. doi:10.1038/nsmb.1727

55. Antos JM, Ingram J, Fang T, Pishesha N, Truttmann MC, Ploegh HL. Site-Specific Protein Labeling via Sortase-Mediated Transpeptidation. Curr Protoc Protein Sci. 2017;89:15.3.1–15.3.19. doi:10.1002/cpps.38

56. Fang T, Duarte JN, Ling J, Li Z, Guzman JS, Ploegh HL. Structurally Defined αMHC-II Nanobody-Drug Conjugates: A Therapeutic and Imaging System for B-Cell Lymphoma. Angew Chem Int Ed Engl. 2016;55(7):2416–2420. doi:10.1002/anie.201509432

57. McColl G, Roberts BR, Pukala TL, et al. Utility of an improved model of amyloid-beta (Aβ1-42) toxicity in Caenorhabditis elegans for drug screening for Alzheimer’s disease. Mol Neurodegeneration. 2012;7(1):57. doi:10.1186/1750-1326-7-57

58. Hunt C, Morimoto RI. Conserved features of eukaryotic hsp70 genes revealed by comparison with the nucleotide sequence of human hsp70. Proc Natl Acad Sci USA. 1985;82(19):6455–6459. doi:10.1073/pnas.82.19.6455

59. Frøkjaer-Jensen C, Davis MW, Hopkins CE, et al. Single-copy insertion of transgenes in Caenorhabditis elegans. Nat Genet. 2008;40(11):1375–1383. doi:10.1038/ng.248

60. Mao S, Qi Y, Zhu H, Huang X, Zou Y, Chi T. A Tet/Q Hybrid System for Robust and Versatile Control of Transgene Expression in C. elegans. iScience. 2019;11:224–237. doi:10.1016/j.isci.2018.12.023

61. Shin Y, Berry J, Pannucci N, Haataja MP, Toettcher JE, Brangwynne CP. Spatiotemporal Control of Intracellular Phase Transitions Using Light-Activated optoDroplets. Cell. 2017;168(1-2):159–171.e14. doi:10.1016/j.cell.2016.11.054

62. Zhang L, Ward JD, Cheng Z, Dernburg AF. The auxin-inducible degradation (AID) system enables versatile conditional protein depletion in C. elegans. Development. 2015;142(24):4374–4384. doi:10.1242/dev.129635

63. Ashley GE, Duong T, Levenson MT, et al. An expanded auxin-inducible degron toolkit for Caenorhabditis elegans. Genetics. 2021;217(3):iyab006. doi:10.1093/genetics/iyab006

