## Supplemental Tables for "HSP-1-Specific Nanobodies Alter Chaperone Function *in vitro* and *in vivo*"

| Target/Epitope | Manufacturer | Catalog # | Blocking Solution | Dilution Solution | Dilution/ Concentration |
|---|---|---|---|---|---|
| RFP | Chromotek/Proteintech | 6g6 | 5% milk/0.1% TBS-T | 2.5% milk/0.1% TBS-T | 1:1,000 |
| α-tubulin | Developmental Studies Hybridoma Bank | 12G10 | 5% milk/0.1% TBS-T | 2.5% milk/0.1% TBS-T | 0.4 ug/mL |
| HSC70 (HSP-1) | Proteintech | 10654-1-AP | 5% milk/0.1% TBS-T | 2.5% milk/0.1% TBS-T | 1:1,000 |
| HA | Cell Signaling | C29F4 | 5% milk/0.1% TBS-T | 2.5% milk/0.1% TBS-T | 1:1,000 |
| Mouse IgG | Cell Signaling | 7076S | N/A | 0.1% TBS-T | 1:10,000 |
| Rabbit IgG | Cell Signaling | 7074S | N/A | 0.1% TBS-T | 1:10,000 |
| StrepTactin-HRP | Bio-Rad | 161038 | N/A | 0.1% TBS-T | 1:10,000 |

### Supplemental Table 1. Antibody descriptions used in this study.

| Fig. 4A/Fig. S4A | WT | B12 | WT (*hsp-1* RNAi) |
|---|---|---|---|
| **Replicate #1** | | | |
| Number of Animals | 123 | 131 | 136 |
| Median Survival | 14 | 12 | 12 |
| **Replicate #2** | | | |
| Number of Animals | 111 | 126 | 110 |
| Median Survival | 15 | 11 | 11 |
| **Replicate #3** | | | |
| Number of Animals | 119 | 132 | 121 |
| Median Survival | 13 | 11 | 11 |

| Fig. 4B/Fig. S4B | WT | B12 | WT (*hsp-1* RNAi) |
|---|---|---|---|
| **Replicate #1** | | | |
| Number of Animals | 112 | 102 | 74 |
| Median Survival | 12 | 12 | 10 |
| **Replicate #2** | | | |
| Number of Animals | 129 | 125 | 116 |
| Median Survival | 13 | 13 | 11 |
| **Replicate #3** | | | |
| Number of Animals | 108 | 94 | 18 |
| Median Survival | 12 | 12 | 10 |

| Fig. 4C/Fig. 74C | WT | B12 |
|---|---|---|
| **Replicate #1** | | |
| Number of Animals | 110 | 128 |
| Median Survival | 10 | 9 |
| **Replicate #2** | | |
| Number of Animals | 119 | 114 |
| Median Survival | 10 | 10 |
| **Replicate #3** | | |
| Number of Animals | 117 | 117 |
| Median Survival | 9 | 9 |

| Fig S4D | WT | B12 | WT (Induced) | B12 (Induced) | WT (*hsp-1* RNAi) |
|---|---|---|---|---|---|
| Number of Animals | 106 | 72 | 128 | 103 | 115 |
| Median Survival | 20 | 20 | 23 | 23 | 11 |

| Fig S4E | WT | H5 | WT (Induced) | H5 (Induced) | WT (*hsp-1* RNAi) |
|---|---|---|---|---|---|
| Number of Animals | 112 | 97 | 91 | 100 | 99 |
| Median Survival | 20 | 20 | 20 | 20 | 10 |

**Supplemental Table 2. Mild heat stress survival data.**

| Fig. 4D (B12) | WT (15 °C) | B12 (15 °C) | WT *hsp-1* RNAi (15 °C) | WT (25 °C) | B12 (25 °C) | WT *hsp-1* RNAi (25 °C) |
|---|---|---|---|---|---|---|
| **Replica #1** | | | | | | |
| Number of Animals Paralyzed | 26 | 29 | 141 | 143 | 129 | 134 |
| Number of Animals Censored | 107 | 121 | 0 | 0 | 0 | 0 |
| Median Paralysis | Undef. | Undef. | 10 | 10 | 4 | 8 |
| **Replica #2** | | | | | | |
| Number of Animals Paralyzed | 28 | 24 | 115 | 102 | 122 | 105 |
| Number of Animals Censored | 75 | 115 | 0 | 0 | 0 | 0 |
| Median Paralysis | Undef. | Undef. | 11 | 9 | 6 | 9 |
| **Replica #3** | | | | | | |
| Number of Animals Paralyzed | 12 | 10 | 101 | 113 | 124 | 113 |
| Number of Animals Censored | 59 | 80 | 0 | 0 | 0 | 0 |
| Median Paralysis | Undef. | Undef. | 10 | 10 | 4 | 8 |

| Fig. 4E (H5) | WT (15 °C) | H5 (15 °C) | WT *hsp-1* RNAi (15 °C) | WT (25 °C) | H5 (25 °C) | WT *hsp-1* RNAi (25 °C) |
|---|---|---|---|---|---|---|
| **Replica #1** | | | | | | |
| Number of Animals Paralyzed | 24 | 36 | 107 | 100 | 129 | 106 |
| Number of Animals Censored | 78 | 104 | 0 | 0 | 0 | 0 |
| Median Paralysis | Undef. | Undef. | 11 | 11 | 11 | 9 |
| **Replica #2** | | | | | | |
| Number of Animals Paralyzed | 61 | 39 | 103 | 86 | 71 | 51 |
| Number of Animals Censored | 53 | 61 | 0 | 0 | 0 | 0 |
| Median Paralysis | Undef. | Undef. | 11 | 11 | 11 | 11 |
| **Replica #3** | | | | | | |
| Number of Animals Paralyzed | 41 | 32 | 140 | 121 | 112 | 133 |
| Number of Animals Censored | 83 | 73 | 0 | 0 | 0 | 0 |
| Median Paralysis | Undef. | Undef. | 12 | 10 | 10 | 10 |

| Fig. 4F (VHH-7) | WT (15 °C) | VHH7 (15 °C) | WT (25 °C) | VHH7 (25 °C) |
|---|---|---|---|---|
| **Replica #1** | | | | |
| Number of Animals Paralyzed | 26 | 26 | 145 | 147 |
| Number of Animals Censored | 107 | 130 | 0 | 0 |
| Median Paralysis | Undef. | Undef. | 10 | 10 |
| **Replica #2** | | | | |
| Number of Animals Paralyzed | 12 | 24 | 113 | 88 |
| Number of Animals Censored | 59 | 96 | 113 | 88 |
| Median Paralysis | Undef. | Undef. | 10 | 10 |
| **Replica #3** | | | | |
| # Paralyzed | 37 | 35 | 142 | 138 |
| # Censored | 95 | 88 | 0 | 0 |
| Median Paralysis | Undef. | Undef. | 10 | 10 |

### Supplemental Table 3. Paralysis data.
